# Directional integration and pathway enrichment analysis for multi-omics data

**DOI:** 10.1101/2023.09.23.559116

**Authors:** Mykhaylo Slobodyanyuk, Alexander T. Bahcheli, Zoe P. Klein, Masroor Bayati, Lisa J. Strug, Jüri Reimand

**Affiliations:** Computational Biology Program, Ontario Institute for Cancer Research, Toronto, ON, Canada; Department of Medical Biophysics, University of Toronto, Toronto, ON, Canada; Department of Molecular Genetics, University of Toronto, Toronto, ON, Canada; Program in Child Health Evaluative Sciences, the Hospital for Sick Children Research Institute, Toronto, ON, Canada; Division of Biostatistics, Dalla Lana School of Public Health, University of Toronto, Toronto, ON, Canada

## Abstract

Omics techniques generate comprehensive profiles of biomolecules in cells and tissues. However, a holistic understanding of the data requires joint multi-omics analyses that are challenging. Here we present DPM, a data fusion method for combining multiple omics datasets using directionality and significance estimates of genes, transcripts, or proteins. DPM allows users to define how the input datasets are expected to interact directionally, reflecting the initial experimental design or regulatory relationships between the datasets. DPM statistically prioritises genes and pathways that change consistently across the datasets, while penalising those violating the constraints. Joint analyses of transcriptomic, proteomic, DNA methylation, and clinical datasets of cancer samples demonstrate how directional integration identifies genes and pathways modulated across omics datasets, highlights those with inconsistent evidence, and reveals candidate biomarkers with prognostic signals in multiple datasets. DPM is implemented in the ActivePathways method and provides a general framework for testing detailed hypotheses in multi-omics data.

## INTRODUCTION

High-throughput omics technologies enable the systematic mapping of genes, transcripts, proteins, and epigenetic states in cells. While data generation methods advance rapidly, data interpretation remains challenging as genes and proteins do not act alone but instead function in complex molecular pathways and interaction networks. Pathway enrichment analysis identifies characteristic biological processes and pathways in omics data to explain underlying experimental conditions or phenotypes ^1^. A common pathway analysis workflow studies lists of significantly altered or expressed genes detected in omics experiments to identify statistical enrichments of biological processes or molecular pathways from databases such as Gene Ontology (GO) ^2^ or Reactome ^3^. Various established tools such as GSEA ^4^, g:Profiler ^5^, and Enrichr ^6^ are widely used for pathway enrichment analysis in basic and biomedical research.

Combining multiple omics datasets is highly beneficial since each experiment provides complementary biological insights. For instance, transcriptomics and proteomics experiments allow us to measure gene and protein expression, post-translational modifications, and signaling network activity. Genomic and epigenomic methods, on the other hand, help us understand genetic variation and gene regulation. Through joint analysis of these complex datasets, we can prioritise genes and pathways and obtain mechanistic and translational insights that can be experimentally validated. Major comprehensive resources like The Cancer Genome Atlas (TCGA), Encyclopedia of DNA Elements (ENCODE), Genotype-Tissue Expression project (GTEx), Clinical Proteomic Tumor Analysis Consortium (CPTAC), and others offer deep multi-omics profiles of human tissues, disease states, and cancer samples ^7-10^ to enable multi-omics analyses.

Multi-omics data analysis presents unique challenges as omics platforms measure various molecules, include distinct experimental and technical biases, and require specific data processing methods ^11^. Comparing genes, transcripts, and proteins directly across the datasets is therefore problematic. A compelling solution to address this complexity involves mapping of omics signals to a common space of pathways and processes ^1^. One powerful approach involves data fusion of statistical significance estimates, such as P-values, that effectively accounts for platform-specific confounding effects, assuming appropriate statistical analyses have been performed upstream. Several computational methods are available for this type of analysis ^12-18^. Pathway-level methods evaluate pathway enrichments in input omics datasets and integrate these as multi-omics summaries ^13,14^. In contrast, gene-level integration methods prioritise genes or proteins across input datasets and then detect multi-omics pathway enrichments ^15-18^. We recently developed ActivePathways that first quantifies all genes through multi-omics data fusion and then finds enriched pathways and their most characteristic genes and contributions from input datasets ^18^.

Multi-omics analyses often fail to consider fundamental directional dependencies in input datasets. For example, the central dogma suggests that mRNA and protein expression levels of genes should correlate positively. Similarly, DNA methylation of gene promoters is a repressive epigenetic mechanism; therefore, lower gene expression often associates with higher level of DNA methylation. Directional dependencies may be integrated in the experimental design. For example, comparing the omics profiles resulting from gene knockout and overexpression experiments may reveal genes and pathways that are regulated downstream of the two perturbations. While cellular control mechanisms like post-transcriptional or post-translational regulation are likely to confound these broad directional dependencies, direct measurements of these additional effects are often not available for analysis. Nonetheless, considering directional dependencies in multi-omics data analysis allows researchers to test more specific hypotheses, prioritise genes and pathways with greater accuracy, reduce false-positive findings, and gain detailed mechanistic insights from the data. Currently, directional methods designed for multi-omics data analysis are lacking, leaving an opportunity for the development of such approaches to further enhance our understanding of complex biological processes.

Here we propose the computational method DPM for directional integration of genes and pathways across multi-omics datasets. DPM employs user-defined constraints to prioritise significant genes or proteins whose directional changes in the omics datasets comply with these constraints. Simultaneously, DPM penalises genes with significant P-values that have inconsistent directions based on the constraints. The flexibility of these constraints makes our method widely applicable to various statistical merging scenarios and experimental designs. To demonstrate our framework, we conduct three case studies: identifying the downstream targets of an oncogenic lncRNA based on transcriptomic profiles from functional experiments in cancer cells; integrating transcriptomic and proteomic data with patient clinical information for cancer biomarker discovery; and characterising the *IDH1*-mutant subtype of glioma by integrating epigenetic, transcriptomic, and proteomic data. The data fusion method DPM is available in the ActivePathways R package. Researchers can utilise this tool to advance their basic biological and biomedical research by gaining valuable insights from multi-omics datasets.

## RESULTS

### Directional multi-omics data integration for gene prioritisation and pathway analysis

We developed a statistical method for multi-omics data fusion that prioritises genes across multiple omics datasets by integrating their P-values and directional changes such as fold-changes (FC). The method, called DPM (directional P-value merging), implements a user-defined *constraints vector* (CV), which specifies the directional associations between the datasets. For each gene, DPM computes a score based on the P-values and directional changes from the omics datasets such that the genes whose directional changes comply with the CV are prioritised while the genes with conflicting directional changes are penalised. The resulting score is derived from input P-values such that highly significant genes are prioritised or penalised strongly while less-significant P-values contribute less to the scoring. DPM builds on our ActivePathways method ^18^ and is based on our directional extension of the empirical Brown’s P-value merging method ^19,20^. For a given gene, a directionally weighted score *X* is computed across *k* datasets as

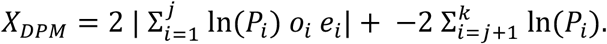

Here, the input P-values *P*_*i*_ and observed directional changes *o*_*i*_ for the gene in dataset *i* are aggregated across two types of datasets: omics datasets (*1 … j*) with directional information and omics datasets *(j+1 … k)* with no directional information, permitting joint analysis of both data types. If either directional or non-directional datasets are not included in the analysis, then the left or right sum in the formula is omitted, respectively. The expected relative dataset direction *e*_*i*_ is obtained from CV. To obtain the merged P-value *P’*_*DPM*_ for the gene reflecting its joint significance in the input datasets given directional information, the scores *X*_*DPM*_ are fit to a cumulative χ^2^ distribution as

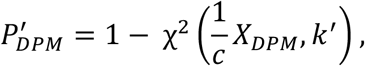

where the degrees of freedom *k’* and scaling factor *c* are estimated from the input P-values empirically ^19^ to account for gene-to-gene covariation for improved significance estimation in omics data with dependencies.

The CV assigns a positive or negative unit sign to each dataset and thereby defines the structure of the multi-omics analysis. Series of positive (+1) or negative (−1) values prioritise genes or proteins in the corresponding datasets that have the same directional changes. In contrast, series of mixed values (+1 and -1) in the CV prioritise genes or proteins that have inverse directional changes in the corresponding datasets. The CV is globally sign-invariant. For example, the CV [+1,+1] for merging two datasets prioritises genes with up-regulation in both datasets or down-regulation in both datasets, and the CV [-1,-1] results in an equivalent analysis. In contrast, the CVs [+1,-1] and [-1,+1] prioritise genes with up-regulation in one dataset and down-regulation in the other dataset, or *vice versa*. The directional changes of genes or proteins from the omics datasets are also only considered as unit signs (*i*.*e*., +1 or -1) because the effect sizes of directions are not directly comparable between omics datasets. Instead, we assume the matching P-values model effect sizes appropriately. Effect size directions may include signs of log-transformed FC values, signs of correlation coefficients, or signs of log-transformed hazard ratio (HR) values from survival analyses. Lastly, the framework permits directionless datasets for which genes or proteins are only prioritised or penalised based on their P-values. For example, mutational burden tests, epigenetic annotations, or network topology analyses often provide P-values but no directional information. Directionless datasets are encoded as zeroes in the CV. In addition to DPM, we also provide directional extensions to Stouffer’s ^21^ and Strube’s ^22^ P-value merging methods based on the METAL method for meta-analysis of genome-wide association studies ^23^. We adapted METAL for joint analyses of directional and non-directional multi-omics datasets (**Methods**).

Our workflow of multi-omics gene prioritisation and pathway enrichment includes four major steps. First, we process upstream omics datasets into a matrix of gene P-values and another matrix of directional values such as FCs (**Figure 1A**). Dedicated upstream processing of the input omics datasets is required to obtain these P-values and FCs. We define a CV with directional constraints based on the overarching hypothesis, the experimental design, and biological insights. We also collect up-to-date pathway information and gene annotations from relevant databases ^2,3^. Second, the matrices of P-values and FCs are merged into a single gene list of P-values using DPM or related methods ^21,22^ (**Figure 1B**). Third, ranked pathway enrichment analysis ^5,18^ is used to statistically associate each pathway to its most enriched fraction of the gene list. It also determines which input omics datasets contribute to each enriched pathway identified in the analysis (**Figure 1C**). Finally, the pathway enrichments are visualised as an enrichment map ^1,24^ that allows users to extract functional themes of biologically related pathways and map their directionality and supporting omics datasets (**Figure 1D**). DPM, combined with pathway enrichment analysis, uses directional biological signals to prioritise genes and pathways across diverse multi-omics datasets.

**Figure 1.**
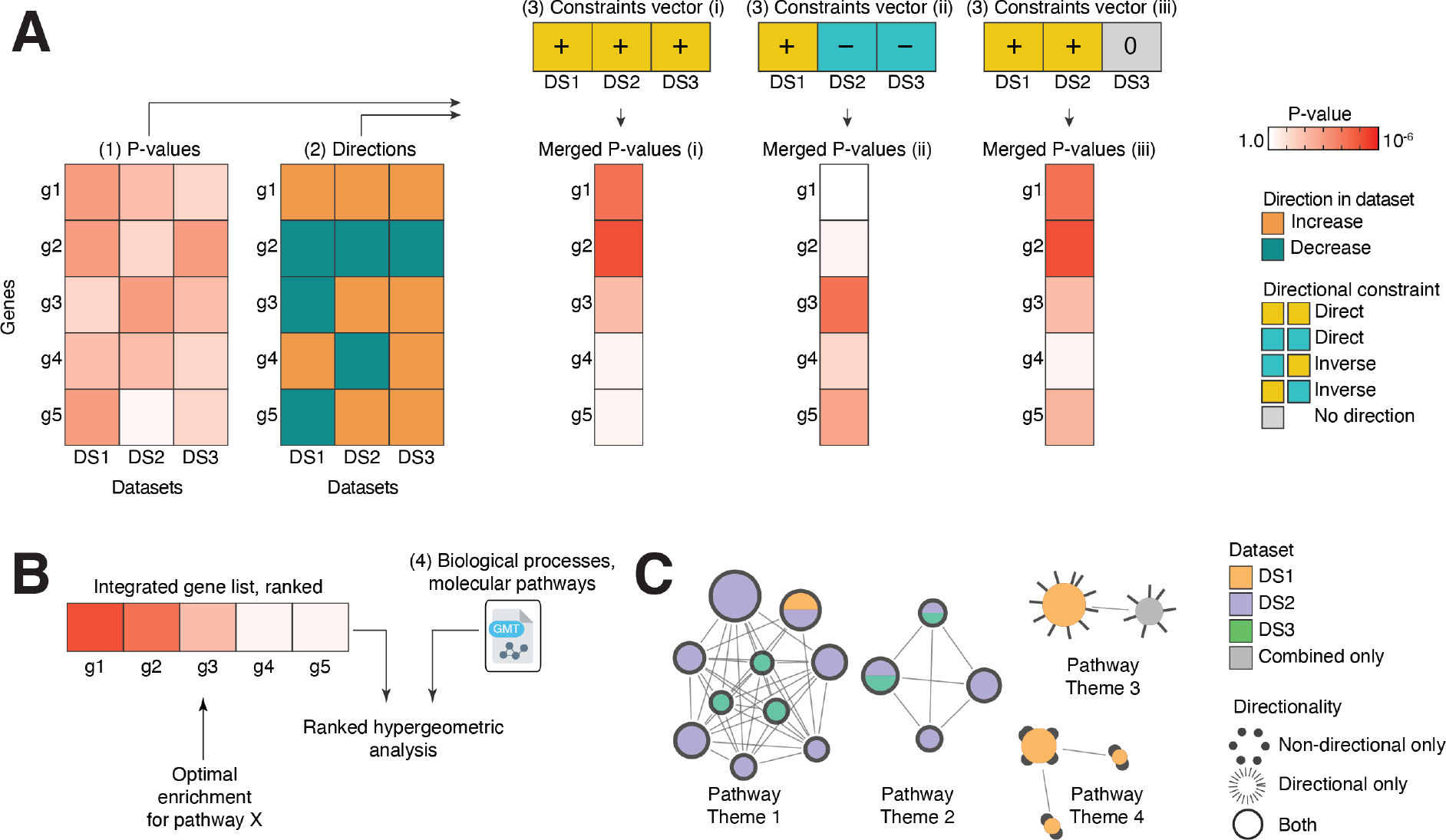
Overview of directional integration of multi-omics data using DPM. **(A)** Four inputs are required: (1) gene activities in multiple input omics datasets quantified as P-values derived upstream (*e*.*g*., differential expression analysis); (2) Directional changes such as fold-change (FC) values or directional coefficients of gene activities, simplified as positive (+1) or negative (−1) unit values. Zeroes are used if no directions are defined in the data; (3) a user-defined constraints vector (CV) of expected directional relationships of the omics datasets; and (4) a file with gene sets of biological processes, pathways, or other functional gene annotations. The DPM method performs data fusion by combining the P-values and directional changes of each gene according to the CV and provides a single integrated gene list of P-values that combines evidence across the input datasets. DPM prioritises genes with significant P-values whose directional changes agree with the CV and penalises genes with disagreements of the CV and the observed directional changes. Three examples of CVs and resulting merged gene lists are shown. **(B)** The integrated gene list is analysed for pathway enrichments using ranked hypergeometric tests in ActivePathways. These tests identify the strongest enrichments in top fractions of the ranked gene list for each pathway. For each pathway, evidence from every input dataset is also evaluated. **(C)** Pathway enrichment results are visualised as an enrichment map that represents a network of enriched pathways where the edges connect pathways with many shared genes. Colours indicate the omics datasets that contribute most to the enrichment, while node outlines indicate whether the pathways were identified using directional or non-directional analyses.

### Benchmarking directional P-value merging

We evaluated our framework by simulating datasets of P-values for 10,000 genes and two experimental conditions. First, we simulated P-values from the uniform distribution that resulted in an expected fraction of significant P-values (*i*.*e*., 5% at P < 0.05), reflecting a dataset with no detectable biological signal. Second, we simulated P-values from an exponential distribution that resulted in an elevated fraction of significant P-values, reflecting a realistic omics dataset with some significantly detected genes or proteins (*i*.*e*., 25% at P < 0.05 or 1% at FDR < 0.05). To model various analysis scenarios, the P-values were then combined into a multi-omics dataset, either as two independent sets of P-values (Pearson r = 1.9 × 10^−4^) or two highly correlated sets of P-values (r = 0.97), reflecting the integration of two completely unrelated or related omics datasets, respectively. We also varied the level of directional agreement of the two input omics datasets and included complete directional agreement, complete directional disagreement, and 50% of directional agreement. We studied the resulting 15 simulated datasets by computing merged P-values using DPM and the modified Strube method and counting the number of nominally significant results at different significance thresholds.

Analysis of simulated P-value datasets revealed the properties of DPM in various multi-omics analysis scenarios (**Figure 2**). When integrating two independent sets of P-values, DPM generated fewer significant results than the modified Strube method. For the negative control scenario of independently generated uniform P-values with full directional agreement, DPM retrieved an expected fraction of significant merged P-values (*i*.*e*., ∼500 at P < 0.05) while two-fold results more were found by the Strube method (**Figure 2A**), suggesting that the latter method may have more false-positive findings than DPM when independent P-values are merged. This was also apparent when integrating exponentially distributed independent P-values and at the 50% level of directional disagreement for correlated and independent P-values (**Figure 2B-C**). As an exception, DPM found more significant results when integrating datasets with full directional disagreement. We studied this in detail by examining the distributions of merged P-values relative to the two sets of input P-values. DPM prioritised genes with directional conflicts if one dataset showed a highly significant P-value while the other dataset only showed limited significance, collectively providing limited significance to the conflicting datasets. In contrast, the Strube’s method assigned more stringent directionality penalties to such genes, suggesting that DPM is more sensitive towards finding genes where the apparent directional disagreement is not supported by statistical significance.

**Figure 2.**
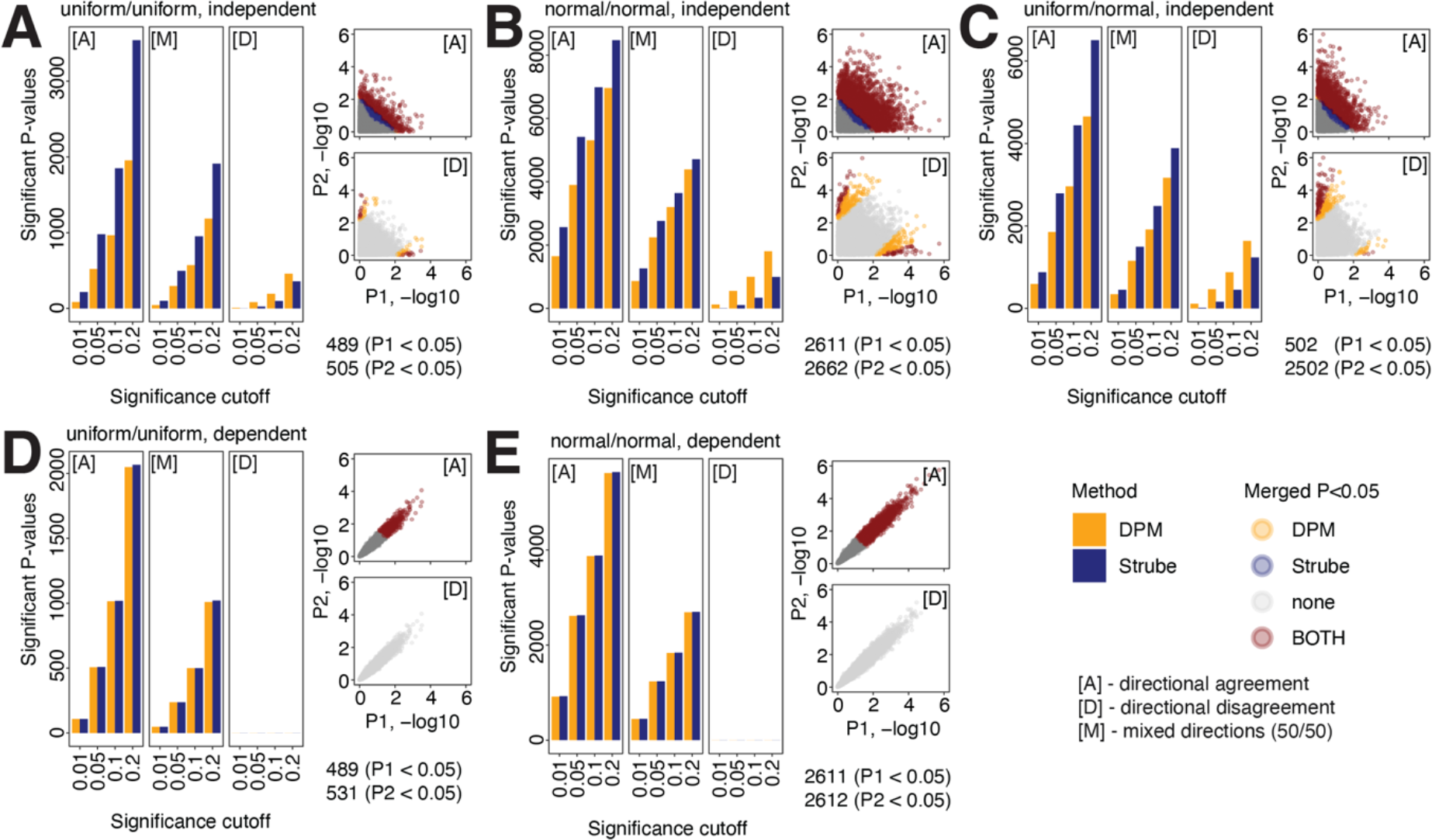
Evaluating directional P-value merging with simulated data. Two sets of 10,000 P-values were simulated using five approaches and were merged using DPM and modified Strube methods. The five approaches are shown in panels A-E. Bar plots on the left show the numbers of significant merged P-values detected at different cut-offs. Scatter plots on the right display the distributions of the two sets of input P-values. Three P-value merging scenarios were considered: all values in directional agreement [A], all values in directional disagreement [D], and mixed directions [M] (*i*.*e*., 50% values in agreement and 50% in disagreement). Numbers of significant input P-values are shown at the bottom right. **(A)** Merging two sets of independent P-values drawn from the uniform distribution. **(B)** Merging two sets of independent P-values drawn from the exponential distribution. **(C)** Merging of independent P-values drawn from uniform and exponential distribution. **(D)** Merging of two sets of correlated P-values drawn from the uniform distribution. **(E)** Merging two sets of correlated P-values drawn the exponential distribution. Scatter plots indicate that DPM is more sensitive to weaker effects seen in a subset of genes with directional conflicts where one of the conflicting datasets is not supported by significant P-values, resulting in a lower penalty.

In contrast to independently generated input P-values, DPM and Strube methods showed very similar performance in merging highly correlated P-values (**Figure 2D-E**). Both methods found the expected fractions of significant merged P-values when integrating the negative control dataset of uniform P-values with full directional agreement. When analysing datasets with full directional disagreements, no significant P-values were found at any of the tested significance thresholds, indicating that both methods applied strong directional penalties were applied to all input P-values. Therefore, directional prioritisation or prioritisation of genes depends on the extent of correlation between the input omics datasets. In summary, this benchmarking exercise demonstrates that directional integration of multi-omics data using DPM is a statistically well-calibrated approach to prioritise or penalise genes via user-defined constraints.

### Integrative analysis of transcriptomic targets of the onco-lncRNA *HOXA10-AS* in glioma

We then studied real omics datasets to evaluate the performance of DPM. First, we analysed our earlier transcriptomics dataset in which the oncogenic lncRNA *HOXA10-AS* was subject to either knockdown (KD) or overexpression (OE) in patient-derived glioblastoma (GBM) cells ^25^. To identify putative direct target genes and pathways of the lncRNA, we used the CV [KD = 1, OE = -1] that prioritised the genes with inverse FC directions in KD and OE experiments and penalised the genes with up-regulation or down-regulation in both experiments (**Figure 3A**). DPM revealed 946 significant genes with the specified directional agreements (FDR < 0.05) (**Figure 3B, Table S1**). On the other hand, we found 640 genes that were significant in the reference non-directional analysis (FDR < 0.05); however, these were penalised when directional constraints were accounted for in DPM. Among prioritised genes, *CPED1* was a top result found by DPM (FDR = 8.2 × 10^−4^) as it was significantly upregulated in the *HOXA10-AS* KD experiment and downregulated in the OE experiment (**Figure 3B**), indicating a potential negative regulatory target of *HOXA10-AS. CPED1* encodes a cadherin and a putative tumor suppressor gene in lung cancer ^29^. The tumor suppressor gene *FAT1* was prioritised due to significant up-regulation in *HOXA10-AS* OE and no significant change in KD, exemplifying another mode of gene prioritisation in DPM. *FAT1* encodes a cadherin protein that is frequently mutated in cancer and contributes to cell proliferation, migration, and invasion ^30,31^, which are hallmarks of advanced glioma. *COL25A1* was a top directionally penalised gene due to significant upregulation in KD and OE experiments (FDR_DPM_ = 0.24, FDR_Brown_ = 1.7x10^−4^) (**Figure 3B**). *COL25A1* encodes a brain-specific membrane-associated collagen protein that binds amyloid beta-peptides ^26^. Other notable directionally penalised genes included *NEGR1*, a neuronal growth regulator, and *CACNA1H*, a calcium voltage-gated channel, that are involved in neuronal development and cell adhesion, respectively ^27,28^.

**Figure 3.**
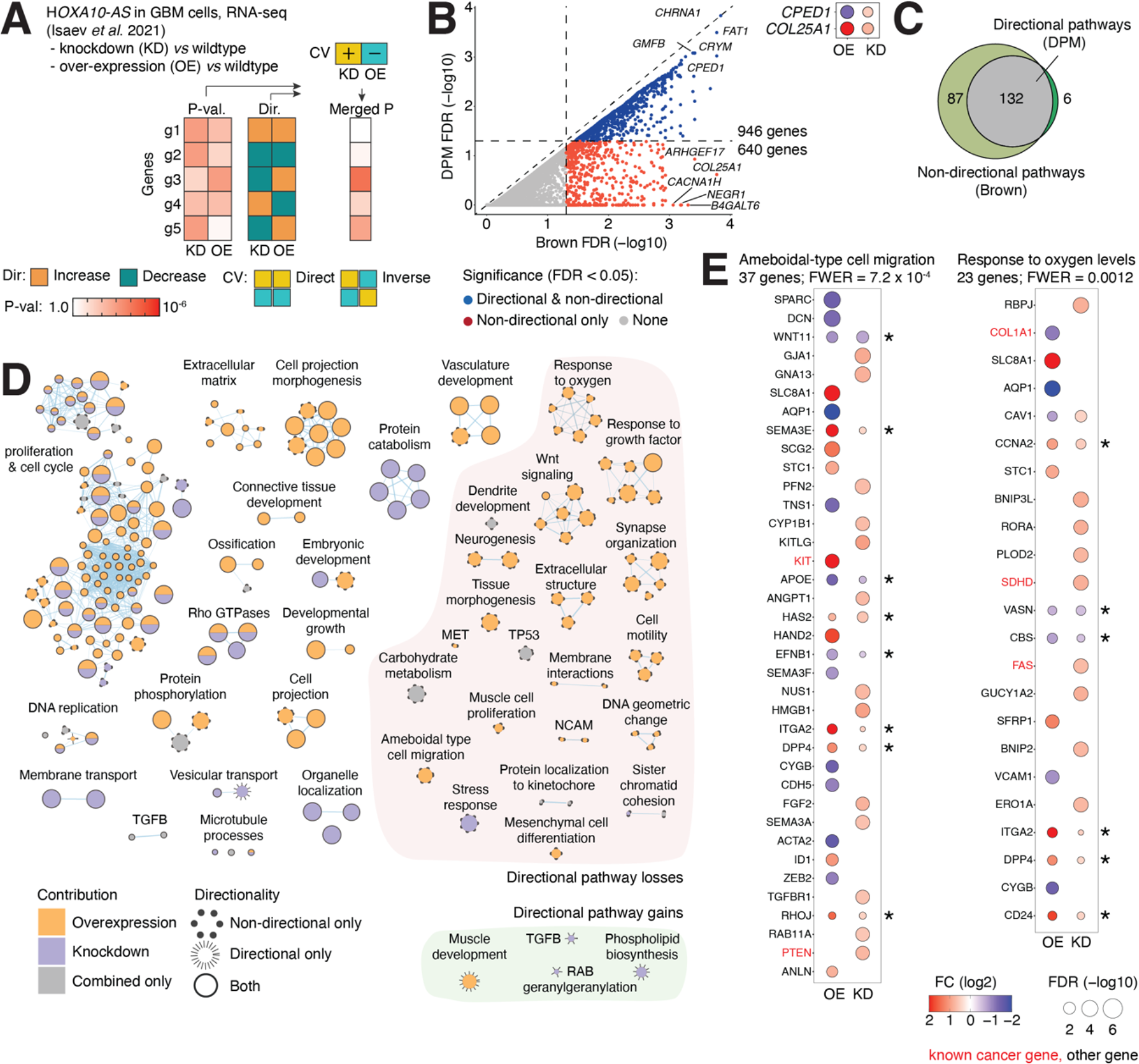
Directional integration of transcriptomics data from functional experiments of *HOXA10-AS* in GBM cells. **(A)** Integrating differential gene expression data from the lncRNA *HOXA10-AS* knockdown (KD) and overexpression (OE) experiments. DPM was configured to prioritise genes that showed different FC directions in KD and OE experiments and penalise the genes with consistent up- or down-regulation in the two experiments. **(B)** Scatter plot comparing integrated gene P-values from DPM (Y-axis) and the non-directional Brown method (X-axis). Significant genes from DPM are shown in blue (FDR < 0.05). Genes with directional agreement are shown along the diagonal while the genes penalised due to directional conflicts appear below the diagonal. Top right: examples of prioritised and penalised genes visualised as FDR and FC values of differential gene expression. **(C)** Venn diagram of enriched pathways found with directional (DPM) and non-directional (Brown) analyses (FWER < 0.05). **(D)** Enrichment map of *HOXA10-AS* target pathways and processes identified in the directional and non-directional analyses (FWER < 0.05). The network shows pathways as nodes that are connected by edges and grouped into subnetworks if the corresponding pathways share many genes. Node colour indicates the dataset contribution (KD, OE, both, or combined-only), and node sizes reflect the number of genes in each pathway. Node outlines show whether the pathways were found using DPM alone (*i*.*e*., directionally prioritised pathway; spiky edges), the non-directional method alone (*i*.*e*., directionally penalised pathways; dotted edges), or were found using both approaches (*i*.*e*., pathways with consistent directions; solid edges). **(E)** GO processes related to cell migration and oxygen levels were penalised in the non-directional analysis due to inconsistent changes in KD and OE conditions. Asterisks indicate genes penalised due to directional conflicts.

Directional pathway analysis using DPM revealed 138 enriched GO processes and Reactome pathways (ActivePathways with DPM, family-wise error rate (FWER) < 0.05) (**Figure 3C-D, Table S2-3)** while the reference non-directional analysis found 219 pathways and processes (ActivePathways with Brown, FWER < 0.05). A third of the enriched pathways from the non-directional analysis (87/219), including cell death, cell motility, brain development, and oxygen response, were excluded by DPM due to directional disagreements in related genes. For example, the GO process of ameboidal-type cell migration found in the non-directional analysis included 37 differentially expressed genes (FWER = 7.3 × 10^−4^). Eight genes showed directional disagreements as these were either upregulated or downregulated in both KD and OE experiments (*WNT11, SEMA3E, APOE, HAS2, EFNB1*, ITGA2, *DPP4, RHOJ*) (**Figure 3E**). Deprioritising these genes using DPM led to the loss of pathway enrichment. Similarly, four oxygen-related processes were lost, such as the GO process describing response to oxygen levels (FWER = 0.0012), in which 6/23 genes had directional disagreements (**Figure 3E**). On the other hand, six pathways were only found by DPM, such as vesicular transport, RAB geranylgeranylation, TGFB signalling, muscle development, DNA replication, and phospholipid biosynthesis, were prioritised through directional information of the pathway genes.

This analysis demonstrates the integration of transcriptomic data from two transcriptomic profiles resulting from opposite functional interventions. Genes and pathways with the expected opposite directional changes in KD and OE experiments may include direct regulatory targets of the *HOXA10-AS* lncRNA that confers phenotypes of advanced glioma ^25^. On the other hand, the penalised genes and pathways with directional disagreements may be regulated indirectly by *HOXA10-AS* through feedback loops or post-transcriptional mechanisms that cannot be measured directly in the omics data we have. However, we can easily prioritise such indirect targets using our method by defining an alternative CV [+1, +1] that selects the genes with matching FCs in KD and OE experiments (**Figure S1**), demonstrating the flexibility of our approach. Integrating the directional associations of omics data from functional experiments improves the resolution of gene prioritisation and pathway enrichment analysis.

### Multi-omics discovery of prognostic biomarkers in transcriptomes and proteomes of ovarian cancer

Next, we integrated cancer transcriptomics and proteomics data from a heterogeneous cancer cohort to associate genes and pathways with patient overall survival (OS) in ten cancer types and 1,140 cancer samples from the CPTAC project ^32,33^ (**Figure 4A, Table S4**). First, we asked which genes significantly associated with OS at the transcript or protein expression level using Cox proportional-hazards (PH) regression with clinical covariates of patient age, sex, and tumor stage. P-values and hazard ratios (HR) for transcript- and protein-level OS associations were integrated using DPM such that genes with consistent OS associations were prioritised while those with inconsistent associations were penalised (*i*.*e*., [RNA = 1, protein = 1]). Ten cancer types were analysed separately (**Figure S3**).

**Figure 4.**
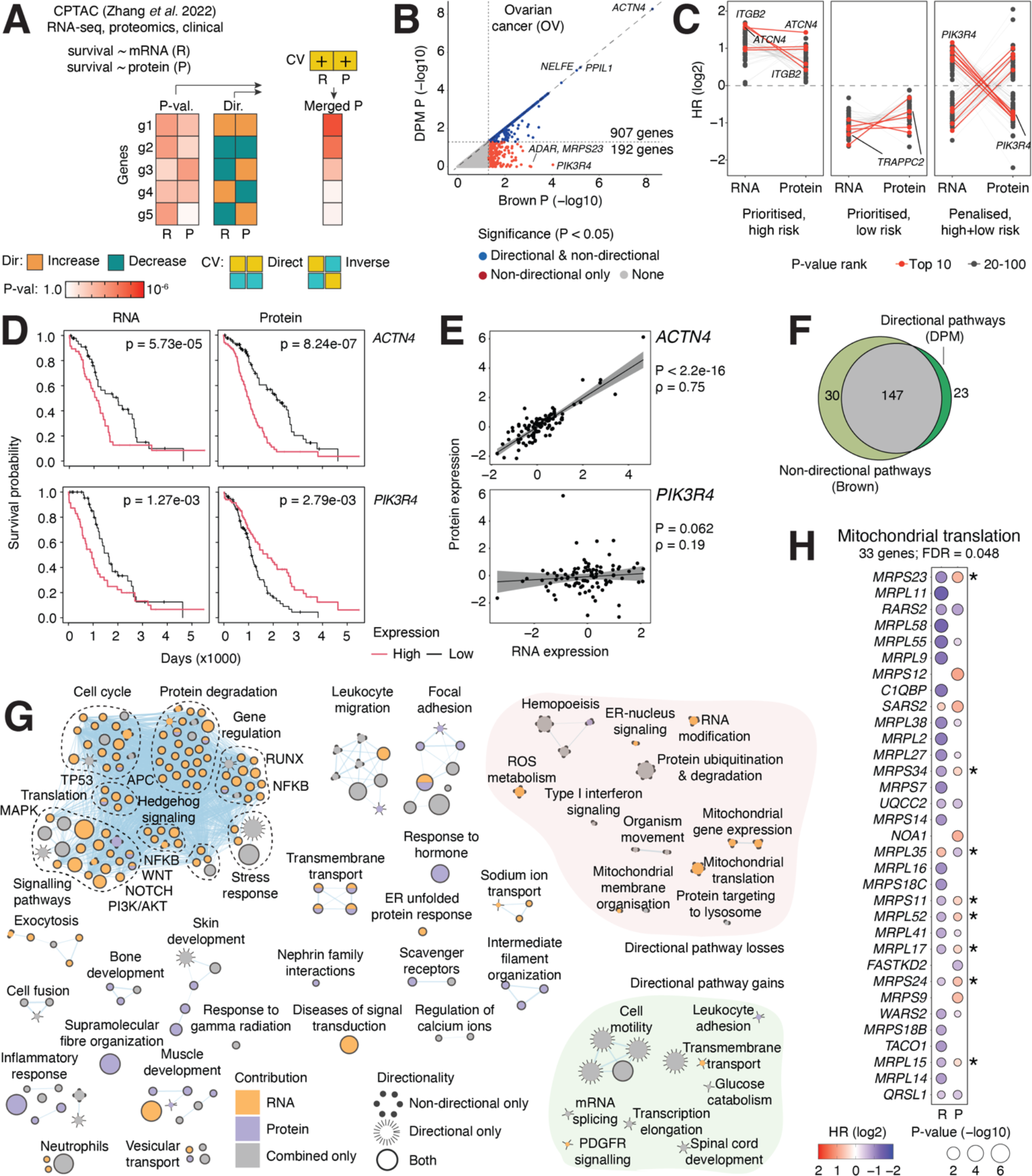
Integrating cancer transcriptomes and proteomes with patient survival information for pathway and biomarker analyses. **(A)** Analysis workflow. mRNA (R) and protein (P) levels for each gene were separately associated with patient overall survival (OS) for ten cancer types in CPTAC using clinical covariates (patient age, patient sex, tumor stage). P-values and hazard ratio (HR) values of mRNA and protein levels retrieved from Cox-PH survival regression models were used for gene prioritisation and pathway analysis. The CV prioritised genes that showed consistent OS associations with transcript and protein levels (*i*.*e*., both positive or both negative) while genes with opposite OS associations were penalised. **(B)** Multi-omics survival associations in ovarian cancer (OV). Directionally prioritised merged P-values of genes from DPM (Y-axis) and non-directional P-values from the reference Brown method (X-axis) are shown. Significant genes from DPM are shown in blue (P < 0.05). Genes along the diagonal have consistent OS associations while the penalised genes with directional disagreements appear below the diagonal. **(C)** Top 100 genes prioritised or penalised by DPM are associated with patient survival with respect to mRNA and protein expression levels and plotted as log-scale HR values. Respective HR values for the same gene are connected by lines. For prioritised genes, both transcript and protein levels associate with higher log-HR (left) or lower log-HR values (middle) reflecting higher or lower patient risk. In, contrast, penalised genes on the right show inconsistent HR values such that lines connecting mRNA- and protein-level associations cross zero. **(D)** Examples of top genes prioritised or penalised by survival associations of mRNA and protein expression in ovarian cancer shown as Kaplan-Meier plots. *ACTN4* (top): high mRNA and high protein levels consistently associate with worse prognosis. *PIK3R4* (bottom): mRNA and protein levels show inconsistent associations with OS. Covariate-adjusted P-values from Cox-PH models and ANOVA are shown. **(E)** Scatterplots of mRNA and protein expression of *ACTN4* and *PIK3R4* in OV explain the OS associations in panel D. Spearman correlation coefficients and P-values are shown. **(F)** Enriched pathways found in genes with OS associations with mRNA and protein levels using directional and non-directional data integration (ActivePathways, FDR < 0.05). Venn diagram shows the pathways prioritised or penalised by directional analysis. **(G)** Enrichment map of pathways and processes with OS associations in transcriptomics and proteomics data in OV (FDR < 0.05). The network shows pathways as nodes that are connected by edges and grouped into subnetworks if the corresponding pathways share many genes. **(H)** The GO process of mitochondrial translation was penalised in the directional analysis due to inconsistent associations. Genes with inconsistent OS associations of mRNA and protein expression are indicated by asterisks.

We focused on the ovarian cancer (OV) cohort with 169 serous cystadenocarcinoma samples. DPM identified 907 genes with consistent survival associations between mRNA and protein levels (P_DPM_ < 0.05) (**Figure 4B, Table S5**). Compared to a reference non-directional analysis, 192 genes were penalised due to inconsistent survival associations (P_Brown_ < 0.05). We examined the survival associations of individual genes to explain the directional integration. Significant genes identified by DPM comprised two groups with either positive or negative OS associations, while the genes penalised by DPM showed both types of associations (**Figure 4C**). *ACTN4*, the most significant prioritised gene (P_DPM_ = 5.4 × 10^−9^), encodes a cytoskeletal actin-binding protein and a well-known oncogene linked to an invasive phenotype and poor prognosis in ovarian cancer ^34,35^. This is confirmed in our analysis: higher transcript and protein expression of *ACTN4* associated with poor prognosis in OV (**Figure 4D**), and *ACTN4* mRNA and protein levels were expectedly highly correlated (Spearman ρ = 0.75, P < 2.2 × 10^−16^) (**Figure 4E**). In contrast, the top penalised gene *PIK3R4* showed inconsistent OS associations: higher transcript expression associated negatively with OS while higher protein expression associated positively, and no significant correlation in transcript and protein expression was apparent (**Figure 4D-E**). *PIK3R4* encodes a regulatory kinase subunit in the PI3K/AKT pathway that regulates cell growth, motility, survival, metabolism, and angiogenesis ^36,37^. Inconsistent expression and survival associations of *PIK3R4* suggest the activity of additional modes of regulation that likely remain masked in these transcriptomics and proteomics datasets.

Pathway analysis with DPM revealed 170 significant pathways and processes with multi-omics survival associations (ActivePathways FDR < 0.05), including major functional themes of proliferation, focal adhesion, cell motility, immune cell activity, development, and signalling pathways such as Hedgehog, Notch, and NFKB (**Figure 4F-G, Table S6-7**). Compared to a reference non-directional analysis, DPM penalised pathways due to directional disagreements in pathway genes in which inverse associations with OS in transcript and protein expression were found. Biological processes of protein translation and degradation, RNA modifications, and mitochondrial function were deprioritised using DPM. This agrees with previous reports that indicated low correlations of transcript and protein expression levels in such genes ^32,38,39^. For example, the non-directional pathway analysis found the enriched process mitochondrial translation, however, it was penalised in the directional analysis with DPM since a large fraction of the pathway genes (8/33) had inconsistent OS associations in transcriptomics and proteomics data (**Figure 4H**). This analysis demonstrates how our directional multi-omics approach can integrate clinical information to discover biomarkers and biological mechanisms in heterogeneous datasets of patient cancer samples.

### Integrating DNA methylation with transcriptomic and proteomics data to dissect molecular phenotypes of *IDH1*-mutant gliomas

Lastly, we integrated DNA methylation, transcriptomics, and proteomics datasets available in TCGA and CPTAC ^33,40^ using an extended design of positive and negative directional associations between the three data modalities. DNA methylation of gene promoters is a known repressive epigenetic mechanism that often correlates with reduced gene expression; therefore, we can obtain more accurate maps of gene and pathway modulation by inversely associating it with transcript and protein expression (**Figure 5A**). We studied this in detail in the TCGA GBM cohort by comparing subsets of glioma samples based on the mutation status of *IDH1. IDH1* encodes isocitrate dehydrogenase 1, a well-established molecular marker of glioma that indicates lower-risk disease ^41^. First, we analysed differential transcript and protein expression and DNA promoter methylation of the molecular phenotype of IDH1-mutant glioma and compared the resulting lists of significant genes. Differential analyses of DNA methylation and transcript expression contributed the most significant genes, perhaps reflecting the hypermethylation phenotype of *IDH1* mutant gliomas ^42^ (**Figure 5B, Table S8**). However, only few genes (32) were found as significant across all three datasets, and the overlaps were even smaller when considering up-regulated and down-regulated genes separately. This highlights opportunities for directional analysis with DPM that combines significance and FC values for gene prioritisation.

**Figure 5.**
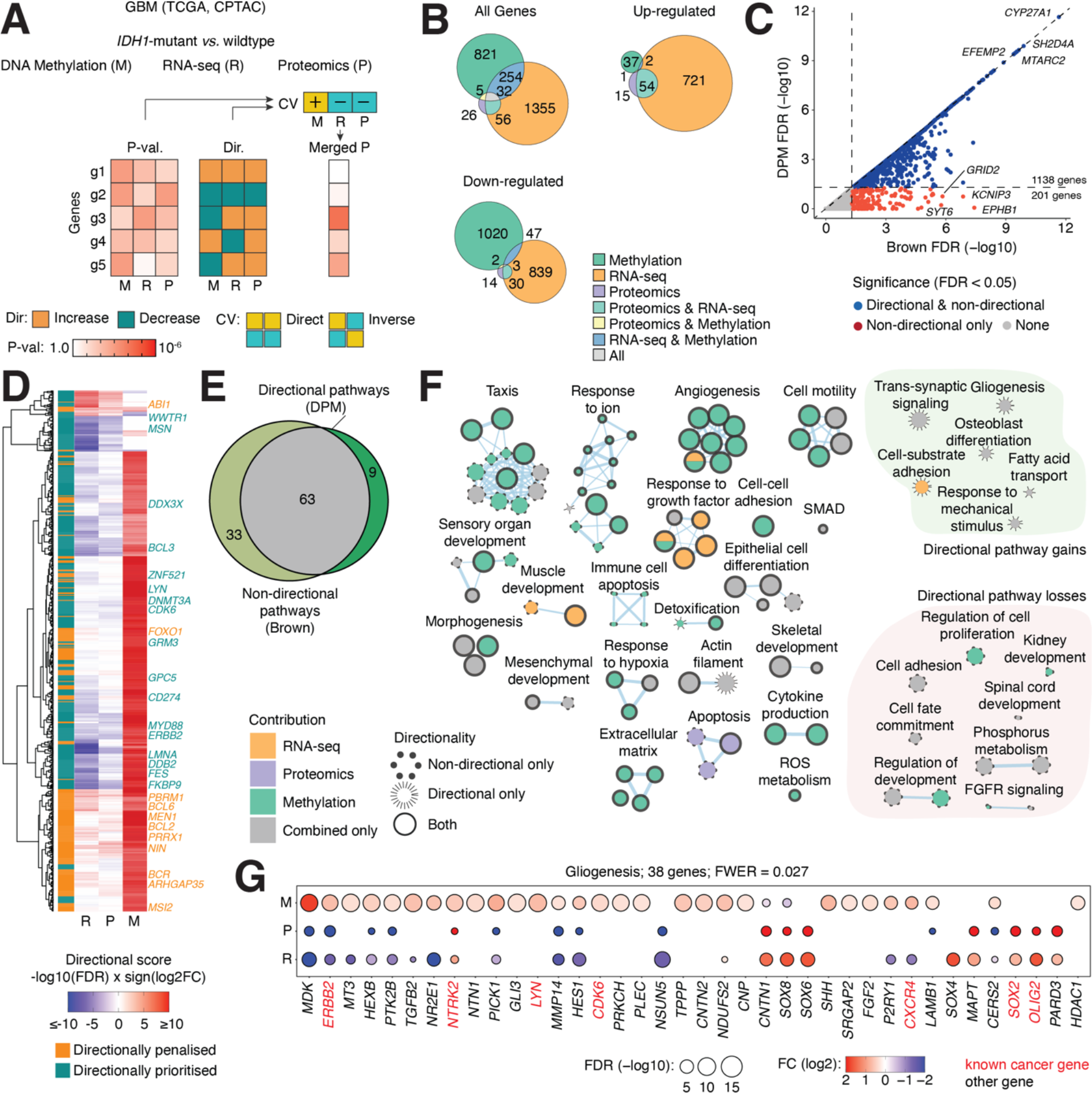
Directional integration of transcriptomics, proteomics, and DNA methylation data to characterise the molecular phenotype of *IDH1*-mutant gliomas. **(A)** Overview of analysis. We compared *IDH1*-wildtype and *IDH1*-mutant gliomas by integrating differential transcript and protein expression and promoter DNA methylation using DPM. The CV defined directional associations between the input datasets: mRNA (R) and protein (P) expression levels associated negatively with DNA promoter methylation (M), as a repressive regulatory mechanism while mRNA and protein levels associated positively with each other. **(B)** Venn diagrams of significant genes found separately in the three datasets (FDR < 0.1). Downregulated genes (bottom left) show reduced mRNA and protein expression and increased promoter methylation, and upregulated genes show decreased promoter methylation and increased expression (top right). **(C)** Scatter plot of directionally prioritised and penalised genes with integrated gene P-values from DPM (Y-axis) and non-directional Brown P-values (X-axis). Significant genes from DPM are shown in blue (FDR < 0.05). Genes with consistent multi-omics signals according to the CV are shown on the diagonal, while the 201 genes below the diagonal have directional disagreements. **(D)** Heatmap of significant genes that were either prioritised or penalised by DPM. The genes were selected stringently using non-directional P-value merging (Brown, FDR < 0.001) and labelled based on DPM as directionally penalised (orange) or prioritised (teal). As expected, prioritised genes were often characterised by high promoter methylation consistently with reduced mRNA and protein expression. Penalised genes often had high promoter methylation and elevated transcript or protein expression that is inconsistent with the CV. Known cancer genes are labelled. **(E)** Venn diagram of enriched pathways from the directional and non-directional analyses (ActivePathways, FWER < 0.05). DPM and Brown methods were used for gene prioritisation, respectively. **(F)** Enrichment map of pathways and processes representing the multi-omics phenotype of *IDH1*-mutant GBM. The network shows pathways as nodes that are connected by edges if the corresponding pathways share many genes. Groups of pathways lost or gained in the directional analysis are grouped on the right. **(G)** The gliogenesis process is significantly detected in the directional analysis and remains undetected in the non-directional analysis. Multiple genes involved in gliogenesis show significant and directionally consistent changes in the three omics datasets that collectively prioritise this process via DPM. Pathway genes with significant multi-omics signals are shown with FDR and FC values.

We performed a directional analysis of the multi-omics dataset by prioritising inverse associations of promoter methylation levels with direct associations of protein and transcript levels using the CV [methylation = +1, mRNA = -1, protein = -1] (**Figure 5A**). This revealed 1138 significant genes (FDR < 0.05, **Figure 5C, Table S8**), while 201 additional genes were penalised due to directional conflicts, compared to the reference non-directional analysis (Brown, FDR < 0.05). The directionally prioritised genes were often driven by high promoter methylation and reduced transcript and protein expression that is consistent with the *IDH1* hypermethylator phenotype. In contrast, the genes penalised by DPM often showed higher promoter methylation combined with upregulation at the transcript or protein level (**Figure 5D**), potentially due to additional post-transcriptional or post-translational regulation that we could not detect reliably. We found 98 known cancer-associated genes using DPM (FDR < 0.05), of which 26 (27%) were consistently regulated between the three datasets. Pathway enrichment analysis of the directionally prioritised genes revealed 72 pathways and processes (FWER < 0.05, ActivePathways, **Table S10**), while 33 pathways identified through a non-directional reference analysis were penalised by DPM (**Figure 5E, Table S9**). DPM penalised biological processes and pathways that appear to be less relevant to glioma biology, such as the muscle organ development process found in the non-directional reference analysis (**Figure 5F**). Many significant genes in the pathway showed directional disagreements (80/195) and were therefore penalised by DPM. Encouragingly, some processes relevant to glioma biology were only found in the directional analysis, such as the process of gliogenesis that defines *IDH1-*mutant gliomas ^43^ (FWER = 0.0207) (**Figure 5G**). As expected, several genes involved in gliogenesis showed significant and directionally consistent changes in *IDH1*-mutant gliomas. For example, the transcription factor *OLIG2* that regulates glial fate and gliomagenesis ^45^ was upregulated in *IDH1*-mutant gliomas at the mRNA and protein level, while the oncogenic receptor tyrosine kinase *ERBB2* that associates with cell survival and proliferation in various cancer types ^44^ was inhibited through the three data modalities. In summary, this case study demonstrates the use of DPM in analysing complex multi-omics datasets for fundamental and translational insights.

## DISCUSSION

We describe a data fusion algorithm that applies user-defined constraints for directional gene prioritisation and pathway enrichment analysis in multi-omics datasets. The method is broadly applicable to various analytical workflows and experimental designs as it relies only on appropriately derived P-values and directional information for all genes. Further, datasets with and without directional information can be analysed jointly. We demonstrate our method by analysing multi-omics datasets of experimental systems and heterogeneous patient cohorts where we encode various directional constraints to capture direct and inverse associations of genes and proteins and pathways. We can also integrate patient clinical information to enable discovery of candidate biomarkers and explore the molecular phenotypes of high-risk disease. A notable limitation of our approach is that directional constraints only provide simplified representation of cellular logic. For example, transcript and protein levels are not always correlated due to additional control mechanisms such as post-translational modifications, protein-protein interactions, alternative splicing, or feedback loops, for which comprehensive molecular data are often not available. Limited transcript-protein correlations have been described in protein translation, mRNA splicing, oxidative phosphorylation, electron transport chain, and other housekeeping processes ^32,38,39,46^. However, our method remains valid given the underlying assumptions. Inverted directional constraints can be used provide further insights: for example, one can map genes and pathways whose transcript and protein levels are inversely associated to study their additional control mechanisms. Thus, the directional constraints provide a useful tool for more accurate hypothesis testing in integrative multi-omics analyses.

Our generic framework is broadly applicable as it makes only a few assumptions about input data. First, accurate upstream data processing is essential for directional multi-omics analyses. Different omics platforms require dedicated preprocessing methods to identify statistically significant signals and account for intrinsic biases. Second, our method relies on accurately computed P-values, which need to be well calibrated and comparable between the input datasets. Third, we only use discrete directional information to reflect increases or decreases in gene or protein activity. Examples include signs of log-transformed fold-changes from differential expression analyses, coefficients from correlation or regression analyses, and hazard ratios from survival analyses. We use discrete directional information as a simple and robust approach that can be adapted to various designs such as case-control comparisons, time series, and cluster analysis and we assume that P-values reflect the strength directional information appropriately. In contrast, numeric directional values would be error-prone as effect sizes of various omics platforms are not comparable directly. Fourth, genes, proteins, transcripts, sites in non-coding DNA, and other elements measured in multi-omics datasets need to be mapped to a common namespace of genes, requiring additional work and compromises in dataset annotation. Lastly, we envision several areas of future work. Our current method is designed for analysing bulk omics datasets and single-cell datasets in common workflows that integrate across a relatively small number of omics profiles or clusters. More work is needed to ensure the scalability of our method to large numbers of multi-omics profiles. Second, our pathway analysis currently uses a simplified representation of molecular pathways and biological processes collapsed into gene sets, however, future data fusion approaches designed for molecular interaction networks can provide complementary insights to gene function and interactions in multi-omics data. In summary, directional multi-omics analysis for gene prioritisation and pathway analysis enables mechanistic and translational insights by focusing on understudied intersections of complex omics datasets.

## METHODS

### Directional P-value merging (DPM)

To integrate multiple omics datasets through gene P-values and directional information, we implemented or repurposed directional extensions to four P-value merging strategies: the methods by Fisher, Brown, Stouffer, and Strube. The methods by Brown and Strube were extended based on the methods by Fisher and Stouffer, respectively, to account for the covariation of gene P-values across input datasets. All methods assume that the P-values are uniformly distributed under the null hypothesis and are well calibrated. Covariation-adjusted methods account for dependencies in the P-value distributions and thereby provide more conservative merged P-values. As omics datasets include biological dependencies, covariation-adjusted methods are usually more appropriate for this type of analysis.

The Fisher’s method for merging P-values ^47,48^ assumes independent P-values are used as input. It collapses *k* P-values *P*_*i*_ to a score *X*_*F*_ that is a sum of log-transformed P-values. The score *X*_*F*_ is transformed into a merged P-value *P’*_*F*_ through the cumulative χ^2^ distribution with 2*k* degrees of freedom:

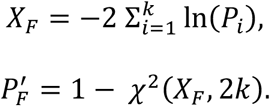

The Brown’s method ^20^ extends the Fisher’s method to account for P-value covariation in input datasets by approximating the score *X*_*F*_ from the Fisher’s method using a scaled *χ*^2^ distribution.

The scaling factor *c* and the updated degrees of freedom *k*^′^ are derived as 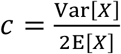 and 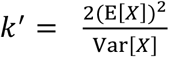, respectively. The expected value and variance of the scaled distribution are derived as E[*cχ*^2^(*k*^′^)] = *ck*^′^ and Var[*cχ*^2^(*k*^′^)] = 2*c*^2^*k*^′^, respectively. The merged Brown P-value *P’*_*B*_ is computed as a sum of log-transformed P-values from the cumulative scaled χ^2^ distribution with the scaling factor *c* and degrees of freedom *k’*, as

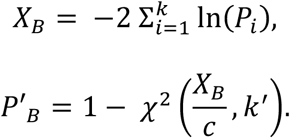

The empirical Brown’s method (EBM) estimates the expected value and variance from the input datasets non-parametrically ^19^. We used EBM here and refer to it as Brown’s method.

To incorporate directionality to the Fisher’s method, we jointly analyse the directional information with the observed gene direction *o*_*i*_ and the expected gene direction *e*_*i*_ in each dataset *i*. For example, in differential gene expression analyses of two conditions relative to a control condition, *o*_*i*_ would be the sign of the fold-change of a gene in condition *i*, and *e*_*i*_ would be the expected relative directional agreement of the two conditions. Both *o*_*i*_ and *e*_*i*_ adopt the values of +1, -1 and 0. The constraint vector (CV) [+1, +1] prioritises genes with consistent fold-change directions across two conditions and is functionally equivalent to the CV [-1, -1] in our method. Alternatively, the CV [+1, -1] or the CV [-1, +1] can be used interchangeably to prioritise genes with opposite fold-change directions across two conditions. Values of zero are used for both *o*_*i*_ and *e*_*i*_ to define datasets where the user intends to not encode directional information, for example acquiring P-values from a gene mutational burden test. The directional coefficients are incorporated in P-value merging to sum log-transformed P-values, as

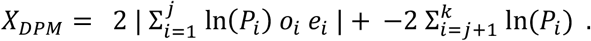

Here, the datasets (1, 2, …, *j*) have defined directional information available while the datasets (*j*+1, *j*+2, …, *k*) do not. This approach permits analyses of mixed directional and non-directional datasets. If either directional or non-directional datasets are not included in the analysis, then the left or right sum is omitted, respectively. Intuitively, directional agreements increase the sums of log-transformed P-values that lead to increased significance of the merged P-value, while directional disagreements reduce the sums. The absolute function is used to ensure that the CV is globally sign invariant (*i*.*e*., [-1,1] ≡ [1,-1] and [1,1] ≡ [-1,-1]). An example is shown in **Figure S2**. Finally, a scaled cumulative *χ*^2^ distribution is computed from Brown’s method to obtain the merged P-values directionally as

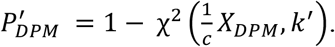

This method is referred to as DPM (directional P-value merging) and is used throughout our study.

In addition to the above, we implemented a directional extension of the METAL method ^49^ that extends Stouffer’s method ^21^ for meta-analysis of GWAS studies. Each study has a direction of effect that reflects the impact each allele has on the observed phenotype. This observed directional term, *o*_*i*_, can either be positive (+1), reflecting an increase in the observed phenotype, or negative (−1), reflecting a decrease. Directional Stouffer’s method introduced by METAL converts P-values from *k* independent tests into Z-scores using the inverse of the standard normal cumulative distribution function Φ^−1^ as

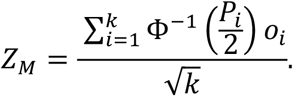

The merged P-values are generated through the standard normal cumulative distribution function, as 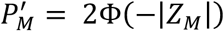. To account for P-value dependencies, Strube’s extension to Stouffer’s method ^22^ leads to more conservative significance estimates by incorporating the overall covariation of P-values in input datasets ^22^, similarly to Brown’s extension of the Fisher’s method. We implemented a directional extension of Strube’s and Stouffer’s methods similarly to METAL as

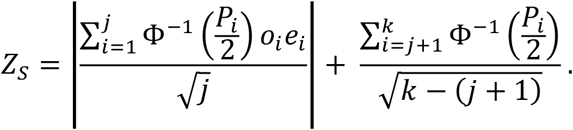

Here, Z scores are acquired for the directional datasets (1, 2, …, *j*) separately from the non-directional datasets (*j*+1, *j*+2, …, *k*) and then each term is combined before calculating a merged P-value, similarly to DPM above.

DPM is available as part of the ActivePathways R package in the CRAN repository (https://cran.r-project.org/web/packages/ActivePathways/index.html).

### Evaluating DPM using simulated and real datasets

We compared DPM and the modified Strube’s method using simulated datasets. The simulated datasets were constructed by generating two sets of 10,000 randomly sampled P-values. First, we created two sets of input P-values independently of each other (IND). Uniformly distributed P-values P_U_ were generated by sampling Z-scores from the normal distribution (μ = 0, σ = 1) and transforming these to P-values relative to the normal distribution (μ = 0, σ = 1). Exponentially distributed P-values P_E_ were generated by sampling Z-scores from the normal distribution (μ = 1, σ = 1) and transforming these to P-values relative to (μ = 1, σ = 1), resulting in an exponential-like distribution that was over-represented in significant P-values (*i*.*e*., ∼25% with P < 0.05). Second, we generated the two sets of input P-values such that the P-values were positively correlated with each other (COR), by first creating one set of Z-scores as described above (*i*.*e*., representing either P_U_ or P_E_) and then adding normally distributed noise (μ = 1, σ = 0.2) to these Z-scores prior to P-value transformation to obtain the second, correlated set of P-values. Spearman correlations of the two sets of P-values were computed. In total, five simulated datasets of P-values were generated: IND(P_U,_ P_U_), IND(P_E,_ P_E_), COR(P_U,_ P_U_), COR(P_E,_ P_E_), and IND(P_U,_ P_E_). We then merged the simulated P-values with directional information in three different configurations: all P-values having directional agreement with the constraints vector, all P-values having directional disagreement, and half of P-values having directional disagreement and half having directional agreement. In the latter case, directional disagreement was assigned randomly using the binomial distribution. Using the resulting 15 configurations of simulated data, we performed directional merging of P-values and counted the numbers of nominally significant merged P-values from DPM and modified Strube methods at different significance thresholds (P < (0.2, 0.1, 0.05, 0.01)).

### Integration of transcriptomics datasets from functional experiments of the HOXA10-AS lncRNA in GBM cells

We analysed the genes and pathways prioritised by directional integration of transcriptomic data from *HOXA10-AS* knockdown (KD) and overexpression (OE) experiments in GBM cells from our earlier study ^25^. We used the CV [KD = -1, OE = 1] to prioritise genes with opposite FCs in the two experiments to account for the inverse modulation of the *HOXA10-AS* lncRNA in the knockout and overexpression experiments. DPM was compared to the non-directional analysis using Brown’s P-value merging. For DPM, we used gene FDR values and FC values for 12,996 protein-coding genes from the original study that filtered previously to exclude very lowly expressed genes. Gene sets of biological processes of Gene Ontology (GO) ^2^ and molecular pathways of Reactome ^3^ were downloaded from the g:Profiler website ^50^ on March 27, 2023. We limited the analysis to gene sets of 10 to 750 genes. The statistical background set included all protein-coding genes. Statistically significant pathways were selected after the default multiple testing correction in ActivePathways (FWER < 0.05). Significantly enriched pathways from the directional and non-directional analyses were merged and visualised as an enrichment map ^24^ in Cytoscape using standard protocols ^1^. Subnetworks were manually organised as functional themes of related pathways. Significant genes in individual pathways were visualised as dot plots with FC and FDR values and cancer genes of the COSMIC Cancer Gene Census database ^51^ were highlighted separately.

### Integration of survival information with transcriptomics and proteomics data in CPTAC

We integrated quantitative proteomic and transcriptomic data of cancer samples with patient survival information obtained from the CPTAC project release 3 ^10^ and TCGA PanCanAtlas dataset ^7^ that included 1,140 cancer samples of ten cancer types: pancreatic, ovarian, colorectal, breast, kidney, head & neck, and endometrial cancer, two subtypes of lung cancer, and brain glioblastoma (**Table S4**). We used the combined dataset assembled by Zhang *et al*. (2022) ^32^ that included transcriptomics data for 15,424 genes and proteomics data for approximately 10,000 genes that varied between cancer types. We used previously processed transcriptomics and proteomics datasets in which transcripts and proteins were measured as standard deviations from median values in the cohorts ^32^. First, we derived directional information from transcript or protein associations with overall survival (OS) based on median dichotomisation of transcript or protein expression. Two Cox proportional-hazards (PH) regression models H0 and H1 were used separately for transcript and protein levels for each gene and in each cancer type. The null Cox-PH model H0 only included clinical covariates as predictors of OS. The alternative Cox-PH model H1 used transcript or protein expression level together with common clinical covariates (patient age, patient sex, tumor stage) as predictors of OS. ANOVA analysis comparing the fits of the models H0 and H1 using a chi-square test was conducted to derive P-values and HR values reflecting transcript- and protein-level OS associations. Second, the directional integration with DPM was conducted using a matrix of transcript and protein P-values from the ANOVA analyses and as directional information the corresponding log-transformed HR values were used.

A non-directional analysis was conducted using the Brown’s method as reference. To handle missing values in the data, genes that had fewer than 20 patients with transcriptomic, proteomic, or clinical information were not analysed and were assigned insignificant values (P = 1, log2HR = 0) in the final input matrices. The CV [RNA = +1, protein = +1] was used to prioritise the genes for which transcript and protein levels associate with OS either positively or negatively, while the genes showing a positive OS association with transcript and a negative association with protein expression (or *vice versa*) should be penalised. Integrative pathway enrichment analysis was performed in the ovarian cancer (OV) dataset similarly to the *HOXA10-AS* dataset described above. We compared the pathway enrichment results between the gene lists prioritised by DPM and as reference, the gene lists prioritised using the non-directional Brown’s method. The background set for pathway analysis included 9,064 genes for which both transcriptomic and proteomic measurements were available. Significant pathways were selected using the more sensitive FDR correction (FDR < 0.05) instead of the default correction Holm FWER method in ActivePathways to account for reduced statistical power of OS associations in heterogeneous datasets of cancer patients.

### Integrative analysis of IDH1-mutant GBMs using transcriptomics, proteomics, and DNA methylation data

We integrated three data modalities with multi-directional constraints: transcriptomics (RNA-seq), quantitative proteomics (isobaric label quantitation analysis with orbitrap), and DNA methylation (CpG Illumina 450k microarray). We studied genes and pathways differentially regulated in a subset of gliomas categorised as glioblastomas (GBMs) that carry a specific missense mutation (R132H) in the *IDH1* gene, a prognostic marker of lower-risk gliomas. We included transcriptomics and DNA methylation datasets from TCGA ^52^ and proteomics data from CPTAC-3 ^53^. GBMs with *IDH1* R132H mutations were identified from the Genomic Data Commons (GDC) web portal using their TCGA patient IDs ^54^. First, we performed differential analyses of transcriptomics, methylation, and proteomics datasets by comparing subsets of GBMs based on their *IDH1* mutation status. We limited the analyses to 10,902 genes for which all three data types were available. Transcriptomics data were downloaded as gene read counts of transcripts per million (TPM) values using the TCGAbiolinks R package ^55^ (May 9th, 2023). We compared the transcriptomes of 7 *IDH1*-mutant (*IDH1* R132H) GBMs and 166 IDH-wildtype GBMs. One GBM sample with a different *IDH1* mutation (R132G) was excluded from all analyses. A differential gene expression analysis of *IDH1*-mutant *vs*. wildtype GBMs was performed non-parametrically using Mann-Whitney U-tests. The resulting P-values for genes were corrected for multiple testing using the Benjamini-Hochberg FDR method. DNA methylation data were downloaded using TCGAbiolinks ^55^ for 6 *IDH1*-mutant GBMs and 149 *IDH1*-wildtype GBMs as beta values measuring CpG site methylation.

We limited the analysis to CpGs in gene promoters using Human EpicV2 annotations ^56^. For each gene, we calculated the mean beta value across the CpG probes in its promoter and conducted a differential methylation analysis of the mean values in *IDH1*-mutant *vs. IDH1*-wildtype GBMs using Mann-Whitney U-tests. P-values were corrected for multiple testing using FDR. Genes with significant but small fold-changes in differential methylation (absolute log2-FC < 0.25) were soft-filtered by assigning insignificant P-values (P = 1). Proteomics data for GBMs was retrieved from the CPTAC-3 project and the dataset processed by Zhang *et al*. (2022)^32^. GBMs carrying *IDH1* R132H mutations were identified in GDC using CPTAC-3 IDs ^54^.

Significant proteome-wide differences in 6 *IDH1*-mutant GBMs (*IDH1* R132H) relative to 92 *IDH1*-wildtype GBMs were evaluated using Mann-Whitney U-tests and P-values corrected for multiple testing using FDR. Gene- and pathway-based multi-omics data integration of the *IDH1*-mutant GBM analysis was performed similarly to the analyses above. The P-values from transcriptomic, methylation, and proteomic data were merged using DPM and the Brown method as a reference. Unadjusted P-values and log2-transformed FC values were used for data integration. The CV was defined as [mRNA = -1, protein = -1, methylation = +1] to prioritise genes with positive associations between transcriptomic and proteomic values and negative associations with DNA methylation in promoters, assuming that high promoter methylation is a repressive gene-regulatory signal that inversely associates with gene expression at the transcript and protein level, while transcript expression directly associates with protein expression. An integrative pathway enrichment analysis was performed similarly to the analyses described above. The statistical background set for the pathway analysis included 10,902 genes. Significant pathways were selected using ActivePathways using default thresholds (Holm FWER < 0.05).

Genes with significant differences in the three datasets were studied using hierarchical clustering and visualised as a heatmap. For the heatmap, unadjusted P-values from the three datasets were merged non-directionally using Brown’s method, corrected for multiple testing using FDR, and filtered for significance using a stringent cut-off (FDR < 0.001). Complete hierarchical clustering was performed using a Euclidean distance metric on directional gene scores (*i*.*e*., -log10(FDR) x sign(log2FC)). Using P-value integration from DPM and the non-directional Brown merging, we categorised the selected genes as either showing or lacking directional agreement between the three omics datasets. Known cancer genes from the COSMIC Cancer Gene Census database ^51^ were labelled in the heatmap.

## Acknowledgments

We would like to thank Dr. Shraddha Pai and Dr. Michael M. Hoffman for helpful discussions. This work was supported by the Discovery Grant of the Natural Sciences and Engineering Research Council (NSERC) to J.R., the New Investigator Award of the Terry Fox Research Institute (TFRI) to J.R., the Canadian Institutes of Health Research (CIHR) Project Grant to J.R., and the Investigator Award to J.R. from the Ontario Institute for Cancer Research (OICR). Funding to OICR is provided by the Government of Ontario. M.S. and M.B. were partially supported by Medical Biophysics fellowships from University of Toronto. A.T.B. was partially supported by the Ontario Graduate Scholarship (OGS). Data used in this publication were partially generated by the Clinical Proteomic Tumor Analysis Consortium (NCI/NIH). The results published here are in part based upon data generated by the TCGA Research Network: https://www.cancer.gov/tcga.

## Author contributions

M.S. developed the method and the software package and performed method benchmarking. M.S. and A.T.B. analysed and interpreted the data. Z.P.K. and M.B. contributed to data analysis and interpretation. L.J.S. contributed to method development and benchmarking. M.S., A.T.B., and J.R. wrote the manuscript. J.R. conceptualised and supervised the project and acquired funding. All authors edited and reviewed the manuscript.

## SUPPLEMENTARY MATERIAL

### Supplementary tables

**Table S1**. Differentially expressed genes in patient-derived GBM cells from the *HOXA10-AS* lncRNA knockdown (KD) and overexpression (OE) experiments.

**Table S2**. Non-directional analysis of enriched pathways in *HOXA10-AS* KD and OE experiments using the Brown’s method.

**Table S3**. Directional analysis of enriched pathways in *HOXA10-AS* KD and OE experiments using DPM.

**Table S4**. Cancer samples with matching transcriptomics and proteomics data in the CPTAC and TCGA datasets.

**Table S5**. Associations of protein and transcript expression levels with patient overall survival (OS) in ovarian cancer.

**Table S6**. Non-directional analysis of enriched pathways with OS associations in transcript and protein expression levels in ovarian cancer using the Brown’s method.

**Table S7**. Directional analysis of enriched pathways with OS associations in transcript and protein expression levels in ovarian cancer using DPM.

**Table S8**. Differential protein and transcript expression, and DNA methylation of *IDH1*-mutant gliomas relative to *IDH1*-wildtype gliomas.

**Table S9**. Non-directional pathway enrichments in *IDH1*-mutant gliomas derived using the Brown’s method.

**Table S10**. Directional pathway enrichments in *IDH1*-mutant gliomas derived using DPM.

### Supplementary figures

**Figure S1**. Directional integration of *HOXA10-AS* transcriptomics data that prioritises genes and pathways with matching changes in knockdown (KD) and overexpression (OE) experiments.

**Figure S2**. A minimal example of merging P-values with directional information across three datasets.

**Figure S3**. Integrating transcriptomic and proteomic signals with cancer patient survival information for prognostic biomarker discovery and pathway analysis in 10 cancer types.

## References

1 Reimand, J. et al. Pathway enrichment analysis and visualization of omics data using g:Profiler, GSEA, Cytoscape and EnrichmentMap. Nat. Protoc. 14, 482–517 (2019). 10.1038/s41596-018-0103-9

2 Ashburner, M. et al. Gene Ontology: tool for the unification of biology. Nature Genetics 25, 25–29 (2000). 10.1038/75556

3 Gillespie, M. et al. The reactome pathway knowledgebase 2022. Nucleic Acids Res. 50, D687–D692 (2022). 10.1093/nar/gkab1028

4 Subramanian, A. et al. Gene set enrichment analysis: A knowledge-based approach for interpreting genome-wide expression profiles. Proc. Natl. Acad. Sci. U. S. A. 102, 15545–15550 (2005). 10.1073/pnas.0506580102

5 Reimand, J., Kull, M., Peterson, H., Hansen, J. & Vilo, J. g:Profiler--a web-based toolset for functional profiling of gene lists from large-scale experiments. Nucleic acids research 35, W193–200 (2007). 10.1093/nar/gkm226

6 Kuleshov, M. V. et al. Enrichr: a comprehensive gene set enrichment analysis web server 2016 update. Nucleic Acids Res 44, W90–97 (2016). 10.1093/nar/gkw377

7 Cancer Genome Atlas Research, N. et al. The Cancer Genome Atlas Pan-Cancer analysis project. Nat Genet 45, 1113–1120 (2013). 10.1038/ng.2764

8 Consortium, E. P. An integrated encyclopedia of DNA elements in the human genome. Nature 489, 57–74 (2012). 10.1038/nature11247

9 Consortium, G. T. The Genotype-Tissue Expression (GTEx) project. Nat Genet 45, 580–585 (2013). 10.1038/ng.2653

10 Edwards, N. J. et al. The CPTAC Data Portal: A Resource for Cancer Proteomics Research. J Proteome Res 14, 2707–2713 (2015). 10.1021/pr501254j

11 Subramanian, I., Verma, S., Kumar, S., Jere, A. & Anamika, K. Multi-omics Data Integration, Interpretation, and Its Application. Bioinform. Biol. Insights 14, 24 (2020). 10.1177/1177932219899051

12 Maghsoudi, Z., Nguyen, H., Tavakkoli, A. & Nguyen, T. A comprehensive survey of the approaches for pathway analysis using multi-omics data integration. Brief. Bioinform. 23, 19 (2022). 10.1093/bib/bbac435

13 Canzler, S. & Hackermuller, J. multiGSEA: a GSEA-based pathway enrichment analysis for multi-omics data. BMC Bioinformatics 21, 13 (2020). 10.1186/s12859-020-03910-x

14 Griss, J. et al. ReactomeGSA-Efficient Multi-Omics Comparative Pathway Analysis. Mol. Cell. Proteomics 19, 11 (2020). 10.1074/mcp.TIR120.002155

15 Xia, J. G. et al. INMEX-a web-based tool for integrative meta-analysis of expression data. Nucleic Acids Res. 41, W63–W70 (2013). 10.1093/nar/gkt338

16 Kaspi, A. & Ziemann, M. mitch: multi-contrast pathway enrichment for multi-omics and single-cell profiling data. BMC Genomics 21, 17 (2020). 10.1186/s12864-020-06856-9

17 Shen, K. & Tseng, G. C. Meta-analysis for pathway enrichment analysis when combining multiple genomic studies. Bioinformatics 26, 1316–1323 (2010). 10.1093/bioinformatics/btq148

18 Paczkowska, M. et al. Integrative pathway enrichment analysis of multivariate omics data. Nat. Commun. 11, 16 (2020). 10.1038/s41467-019-13983-9

19 Poole, W., Gibbs, D. L., Shmulevich, I., Bernard, B. & Knijnenburg, T. A. Combining dependent P-values with an empirical adaptation of Brown’s method. Bioinformatics 32, 430–436 (2016). 10.1093/bioinformatics/btw438

20 Brown, M. B. 400: A Method for Combining Non-Independent, One-Sided Tests of Significance. Biometrics 31, 987 (1975). 10.2307/2529826

21 Stouffer, S. A., Edward A. Suchman, Leland C. DeVinney, Shirley A. Star, Robin M. Williams, Jr. Studies in Social Psychology in World War II: The American Soldier. Princeton: Princeton University Press 1 (1949).

22 Strube, M. J. COMBINING AND COMPARING SIGNIFICANCE LEVELS FROM NONINDEPENDENT HYPOTHESIS TESTS. Psychol. Bull. 97, 334–341 (1985). 10.1037/0033-2909.97.2.334

23 Willer, C. J., Li, Y. & Abecasis, G. R. METAL: fast and efficient meta-analysis of genomewide association scans. Bioinformatics 26, 2190–2191 (2010). 10.1093/bioinformatics/btq340

24 Merico, D., Isserlin, R., Stueker, O., Emili, A. & Bader, G. D. Enrichment Map: A Network-Based Method for Gene-Set Enrichment Visualization and Interpretation. PLoS One 5, 12 (2010). 10.1371/journal.pone.0013984

25 Isaev, K. et al. Pan-cancer analysis of non-coding transcripts reveals the prognostic onco-lncRNA HOXA10-AS in gliomas. Cell Reports 37, 26 (2021). 10.1016/j.celrep.2021.109873

26 Tong, Y., Xu, Y., Scearce-Levie, K., Ptacek, L. J. & Fu, Y. H. COL25A1 triggers and promotes Alzheimer’s disease-like pathology in vivo. Neurogenetics 11, 41–52 (2010). 10.1007/s10048-009-0201-5

27 Noh, K. et al. Negr1 controls adult hippocampal neurogenesis and affective behaviors. Mol. Psychiatr. 24, 1189–1205 (2019). 10.1038/s41380-018-0347-3

28 Sheng, L. F., Leshchyns’ka, I. & Sytnyk, V. Neural Cell Adhesion Molecule 2 Promotes the Formation of Filopodia and Neurite Branching by Inducing Submembrane Increases in Ca2+ Levels. J. Neurosci. 35, 1739–1752 (2015). 10.1523/jneurosci.1714-14.2015

29 Hsu, Y. L. et al. Identification of novel gene expression signature in lung adenocarcinoma by using next-generation sequencing data and bioinformatics analysis. Oncotarget 8, 104831–104854 (2017). 10.18632/oncotarget.21022

30 Pastushenko, I. et al. Fat1 deletion promotes hybrid EMT state, tumour stemness and metastasis. Nature 589, 448-+ (2021). 10.1038/s41586-020-03046-1

31 Morris, L. G. T. et al. Recurrent somatic mutation of FAT1 in multiple human cancers leads to aberrant Wnt activation. Nature Genetics 45, 253–261 (2013). 10.1038/ng.2538

32 Zhang, Y. Q., Chen, F. J., Chandrashekar, D. S., Varambally, S. & Creighton, C. J. Proteogenomic characterization of 2002 human cancers reveals pan-cancer molecular subtypes and associated pathways. Nat. Commun. 13, 19 (2022). 10.1038/s41467-022-30342-3

33 Ellis, M. J. et al. Connecting genomic alterations to cancer biology with proteomics: the NCI Clinical Proteomic Tumor Analysis Consortium. Cancer Discov 3, 1108–1112 (2013). 10.1158/2159-8290.CD-13-0219

34 Yamamoto, S. et al. Actinin-4 gene amplification in ovarian cancer: a candidate oncogene associated with poor patient prognosis and tumor chemoresistance. Mod. Pathol. 22, 499–507 (2009). 10.1038/modpathol.2008.234

35 Tentler, D., Lomert, E., Novitskaya, K. & Barlev, N. A. Role of ACTN4 in Tumorigenesis, Metastasis, and EMT. Cells 8, 16 (2019). 10.3390/cells8111427

36 Huang, J. et al. Frequent Genetic Abnormalities of the PI3K/AKT Pathway in Primary Ovarian Cancer Predict Patient Outcome. Gene Chromosomes Cancer 50, 606–618 (2011). 10.1002/gcc.20883

37 Yang, J. et al. Targeting PI3K in cancer: mechanisms and advances in clinical trials. Mol. Cancer 18, 28 (2019). 10.1186/s12943-019-0954-x

38 Zhang, H. et al. Integrated Proteogenomic Characterization of Human High-Grade Serous Ovarian Cancer. Cell 166, 755–765 (2016). 10.1016/j.cell.2016.05.069

39 Clark, D. J. et al. Integrated Proteogenomic Characterization of Clear Cell Renal Cell Carcinoma. Cell 179, 964-+ (2019). 10.1016/j.cell.2019.10.007

40 Liu, J. et al. An Integrated TCGA Pan-Cancer Clinical Data Resource to Drive High-Quality Survival Outcome Analytics. Cell 173, 400–416 e411 (2018). 10.1016/j.cell.2018.02.052

41 Cohen, A. L., Holmen, S. L. & Colman, H. IDH1 and IDH2 mutations in gliomas. Curr Neurol Neurosci Rep 13, 345 (2013). 10.1007/s11910-013-0345-4

42 Bledea, R. et al. Functional and topographic effects on DNA methylation in IDH1/2 mutant cancers. Sci Rep 9, 16830 (2019). 10.1038/s41598-019-53262-7

43 Venneti, S. et al. Histone 3 lysine 9 trimethylation is differentially associated with isocitrate dehydrogenase mutations in oligodendrogliomas and high-grade astrocytomas. J Neuropathol Exp Neurol 72, 298–306 (2013). 10.1097/NEN.0b013e3182898113

44 Yu, D. & Hung, M. C. Overexpression of ErbB2 in cancer and ErbB2-targeting strategies. Oncogene 19, 6115–6121 (2000). 10.1038/sj.onc.1203972

45 Ligon, K. L. et al. Olig2-regulated lineage-restricted pathway controls replication competence in neural stem cells and malignant glioma. Neuron 53, 503–517 (2007). 10.1016/j.neuron.2007.01.009

46 Komili, S. & Silver, P. A. Coupling and coordination in gene expression processes: a systems biology view. Nat. Rev. Genet. 9, 38–48 (2008). 10.1038/nrg2223

47 Fisher, R. A. 224A: Answer to Question 14 on Combining independent tests of significance. The American Statistician 2 (1948).

48 Fisher, R. (Oliver and Boyd, 1932).

49 Willer, C. J., Li, Y. & Abecasis, G. R. METAL: fast and efficient meta-analysis of genomewide association scans. Bioinformatics 26, 2190–2191 (2010). 10.1093/bioinformatics/btq340

50 Raudvere, U. et al. g:Profiler: a web server for functional enrichment analysis and conversions of gene lists (2019 update). Nucleic Acids Res. 47, W191–W198 (2019). 10.1093/nar/gkz369

51 Tate, J. G. et al. COSMIC: the Catalogue Of Somatic Mutations In Cancer. Nucleic Acids Res. 47, D941–D947 (2019). 10.1093/nar/gky1015

52 Weinstein, J. N. et al. The Cancer Genome Atlas Pan-Cancer analysis project. Nature Genetics 45, 1113–1120 (2013). 10.1038/ng.2764

53 Edwards, N. J. et al. The CPTAC Data Portal: A Resource for Cancer Proteomics Research. J. Proteome Res. 14, 2707–2713 (2015). 10.1021/pr501254j

54 Grossman, R. L. et al. Toward a Shared Vision for Cancer Genomic Data. N. Engl. J. Med. 375, 1109–1112 (2016). 10.1056/NEJMp1607591

55 Colaprico, A. et al. TCGAbiolinks: an R/Bioconductor package for integrative analysis of TCGA data. Nucleic Acids Res. 44, 11 (2016). 10.1093/nar/gkv1507

56 Zhou, W. D., Laird, P. W. & Shen, H. Comprehensive characterization, annotation and innovative use of Infinium DNA methylation BeadChip probes. Nucleic Acids Res. 45, 12 (2017). 10.1093/nar/gkw967

