## Supplementary Figures for "Directional integration and pathway enrichment analysis for multi-omics data"

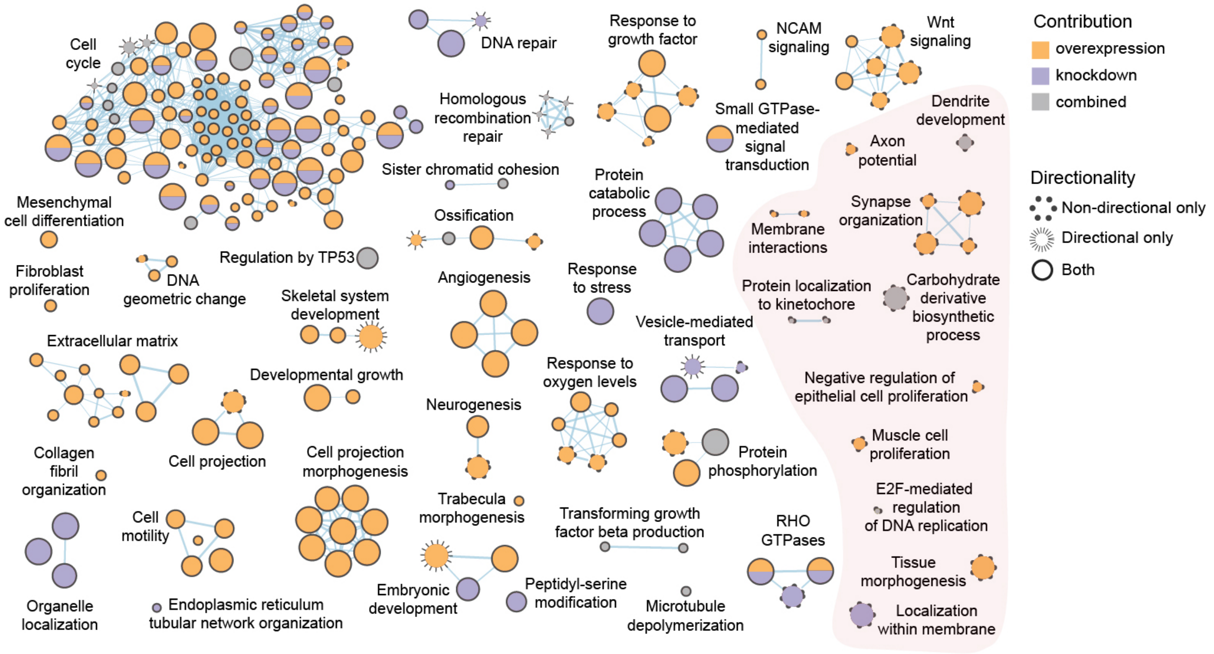


**Figure S1. Directional integration of *HOXA10-AS* transcriptomics data that prioritises genes and pathways with matching changes in knockdown (KD) and overexpression (OE) experiments.** Consistent fold-change (FC) directions for gene prioritisation were encoded in the constraints vector as [KD = +1, OE = +1]. Enrichment map of pathways and processes enriched in the integrative analysis of *HOXA10-AS* KD and OE experiments is shown (ActivePathways, FWER < 0.05). The network includes pathways as nodes that are connected by edges and grouped into subnetworks if the corresponding pathways share many genes. Node color indicates the dataset contribution (KD, OE, both, or combined-only), and node sizes reflect the number of genes per pathway. Node outline indicates whether the pathways were identified using DPM alone (*i.e.*, the directional information helped prioritise pathway genes; spiky edges), directionless Brown alone (*i.e.*, the directional information helped penalise pathway genes with inconsistent fold-changes; dotted edges), or using both approaches (*i.e.*, most pathway genes were consistent with directional information; solid edges). Pink background highlights the major functional themes that were lost in directional analysis.

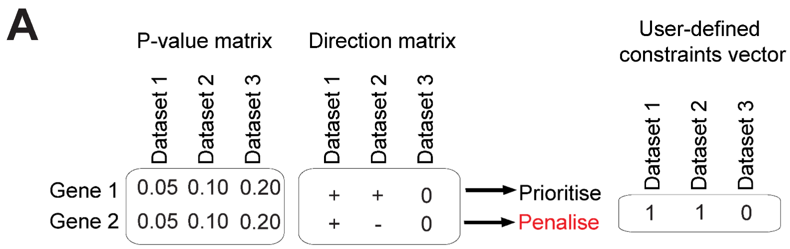


**B**

$$d_i = o_i \, e_i$$

$$X_{\mathrm{DPM}} = 2\left|\sum_{i=1}^{j} d_i \ln(P_i)\right| - 2\sum_{i=j+1}^{k} \ln(P_i)$$

$$X_{\mathrm{DPM}} = 2|d_1 \ln(P_1) + d_2 \ln(P_2)| - 2\ln(P_3)$$

$$P'_{\mathrm{DPM}} = 1 - \chi^2\left(\frac{X_{\mathrm{DPM}}}{c}, k'\right)$$

**Gene 1**

$$d = [1,1,0]\,[1,1,0] = [1,1,0]$$

$$X_{\mathrm{DPM}} = 2|(1)\ln(0.05) + (1)\ln(0.10)| - 2\ln(0.20)$$

$$X_{\mathrm{DPM}} = 13.81$$

$$P'_{\mathrm{DPM}} = 1 - \chi^2\left(\frac{13.81}{1}, 6\right) = 0.03$$

**Gene 2**

$$d = [1,-1,0]\,[1,1,0] = [1,-1,0]$$

$$X_{\mathrm{DPM}} = 2|(1)\ln(0.05) + (-1)\ln(0.10)| - 2\ln(0.20)$$

$$X_{\mathrm{DPM}} = 4.61$$

$$P'_{\mathrm{DPM}} = 1 - \chi^2\left(\frac{4.61}{1}, 6\right) = 0.60$$

**Figure S2. A minimal example of merging P-values with directional information across three datasets. (A)** Three datasets of two genes with identical P-values and differing observed directional information are analysed. Three inputs are required: gene P-values in input omics datasets from upstream analyses; directional coefficients of gene activities, such as fold-change values, simplified as positive (+1) or negative (-1) unit values or zero (0) if no directions are defined; a constraints vector (CV) of expected directional relationships of the omics datasets as defined by the user. Here, the third dataset contains no directional information, while the first two datasets are expected to have a direct (positive) relationship. **(B)** Example of prioritising and penalising genes using directional information. Gene 2 is penalised due to a directional inconsistency while Gene 1 is prioritised. The direction $d_i$ term is acquired by element-wise multiplication of each gene's observed direction ($o_i$) against its expected direction ($e_i$). Chi-square values are acquired for the directional datasets (1 and 2) separately from the non-directional third dataset, and then the magnitude of each term is combined. Unlike Gene 1, Gene 2 does not receive a significant merged P-value due to the observed directional inconsistency.

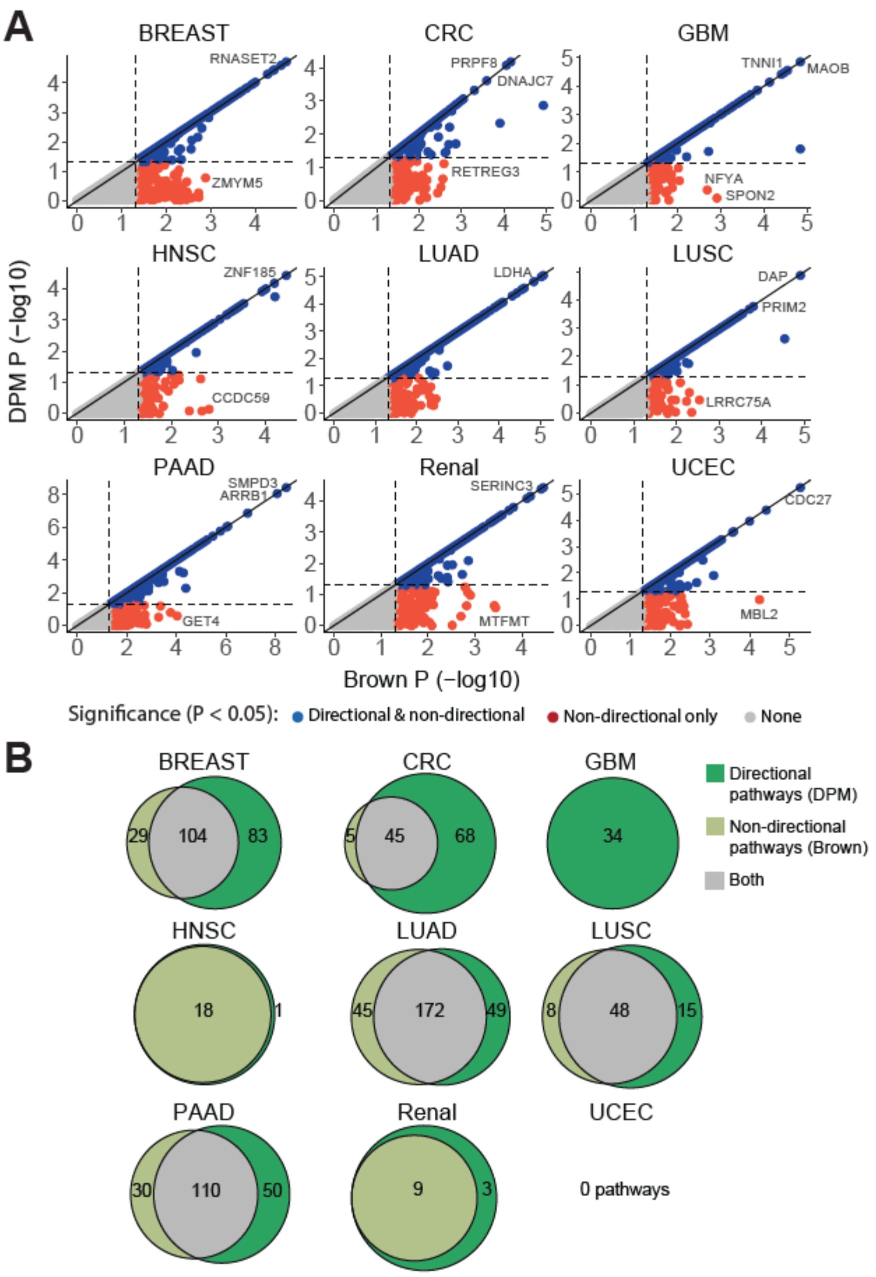


**Figure S3. Integrating transcriptomic and proteomic signals with cancer patient survival information for prognostic biomarker discovery and pathway analysis in 10 cancer types.** Survival analysis was performed using Cox proportional-hazards models with protein or transcript levels and clinical covariates (patient age, sex, and tumor stage) as predictors of overall survival (OS). Each cancer type profiled in the CPTAC and TCGA projects was analysed separately. Each gene was modelled twice: once with transcript level as OS predictor and once with protein level as OS predictor. **(A)** Gene P-values and log2-transformed hazard ratio (HR) values for transcripts and protein levels were integrated. The scatter plots compare integrated P-values of genes from DPM (Y-axis) and non-directional P-value merging (Brown's method; X-axis). Significant genes from DPM are shown in blue ($P < 0.05$). The diagonal shows genes where transcript and protein levels show consistent OS associations, while the genes below the diagonal have directional conflicts with OS between transcript and protein expression levels. **(B)** Pathway-level integration of OS with transcriptomics and proteomics data with and without directional information ($FDR < 0.05$). The Venn diagrams show how many significant pathways were found in the directional and non-directional analyses. DPM and Brown's method were used to prioritise gene lists for pathway analyses.
